# Dynamic brain activity during associative learning examined with MEG/fMRI co-processing

**DOI:** 10.1101/2021.09.10.459821

**Authors:** Sangeeta Nair, Jane B. Allendorfer, Yingying Wang, Jerzy P. Szaflarski

**Affiliations:** University of Alabama at Birmingham, Department of Psychology; University of Nebraska-Lincoln, Department of Special Education and Communication Disorders (SECD); University of Alabama at Birmingham, Department of Neurology

**Keywords:** MEG, fMRI, associative learning, active generation, multiple sparse priors, multimodal data fusion

## Abstract

**Background:** Due to limitations of individual neuroimaging methods we examine spatial and temporal contributions to self-generation using multimodality imaging with functional magnetic resonance imaging (fMRI) and magnetoencephalography (MEG) within the Bayesian framework Multiple Sparse Priors (MSP).

**New Method:** 24 healthy participants performed an fMRI and MEG paired-associate learning task. FMRI data were processed within Group ICA fMRI Toolbox. Independent components (ICs) were temporally sorted by task time series (|r|>0.30 threshold identified task-related ICs). Task-positive (“generate”) ICs were retained as spatial *priors* for MEG analyses. MEG data were processed by averaging trials to increase the signal-to-noise ratio within subjects and with an event-related theta power approach. MEG source reconstructions were constrained within the task-positive ICs for both analytical approaches.

**Results:** For fMRI, five networks were identified as task-related. Four ICs underlying active generation spanned bilateral parietal, orbitofrontal, medial frontal and superior temporal regions, and occipital lobe. FMRI-constrained MEG source reconstructions yielded early visual cortex activity followed by left inferior frontal gyrus (IFG) and orbito-frontal cortex (OFC) recruitment to coalesce in the left inferior temporal lobe. For the event-related theta approach, reconstructions showed a progression of activity from bilateral temporal areas to left OFC and middle temporal gyrus, followed by right IFG.

**Comparison with Existing Methods:** MSP analyses informed by fMRI produced more focused regional activity than reconstructions without *priors* suggesting increased attention and maintenance when selecting relevant semantic information during active generation.

**Conclusions:** Constraining MEG source reconstruction to fMRI *priors* during active generation implicates interconnected fronto-temporal and fronto-parietal networks across time.

## 1. INTRODUCTION

The processes behind acquiring new knowledge are broadly divided into two learning styles: passive and active (Mcdaniel, Waddill, and Einstein 1988; Olofsson and Nilsson 1992; Pugh et al. 1996). Passive learning is conceptualized as a one-way transference of information. Individuals are meant to passively absorb and retain new knowledge through traditional approaches like lectures, presentations, and reading. In contrast, active learning requires engagement and participation from the individual such as group discussions, hands-on workshops, or interactive games (Prince 2004). As these two styles of learning require different contributions from the individual in terms of cognitive mechanisms including attention and depth of processing, they also present specific strengths in various settings. The benefits of active learning have been widely studied across a range of metrics, including improved retention of information, academic achievement, and self-esteem (Mcdaniel et al. 1988; Olofsson and Nilsson 1992). Enhanced memory of active learning, for example, with self-generating content, has been hypothesized to be a result of the increased distinctiveness of the target word (Otten, Henson, and Rugg 2001a).

The neurological processes underlying both passive and active learning recruit distinct and overlapping brain networks. There have been recent efforts to evaluate neuroanatomical and functional contributions to verbal learning in healthy and diseased populations using positron emission tomography (PET) and functional magnetic resonance imaging (fMRI) (Binder et al. 2009), as well as contributions from the electrophysiological literature outlining stereotyped responses across groups (Marinković 2004). However, such approaches are hampered by either the poor temporal resolution of fMRI or poor spatial resolution of PET or electrophysiology techniques. In fMRI, the blood oxygen level dependent (BOLD) responses are sensitive to functional changes in response to a specific task. Although fMRI is often utilized for its high spatial resolution, the method is limited in what information it can provide within the temporal domain. As neuronal activity takes place in the order of milliseconds, considerably faster than any associated vascular changes at roughly 5-7 seconds of the hemodynamic response function (HRF) (Hawco et al. 2007; de Munck et al. 2007), fMRI is unable to preserve the temporal resolution of neuronal activity and thus is not widely used to explore the timing and information flow of cognitive processes. Previously mentioned fMRI studies of healthy controls identify distributed networks that underlie various verbal memory processes (encoding and retrieval), but little is known about the dynamic communication between nodes in these networks (Baker et al. 2001; Buckner, Kelley, and Petersen 1999; Kim 2011).

Bioelectric-based methods, including electroencephalography (EEG) and magnetoencephalography (MEG), are sensitive to electromagnetic fields generated by synaptic currents in the brain. These techniques offer insight into neuronal activity with an especially high temporal resolution, allowing for evaluation of activity as it unfolds across milliseconds. As such, bioelectric and electromagnetic techniques are particularly well suited to explore questions around the dynamics of brain activity and information transfer. While both EEG and MEG noninvasively measure electromagnetic fields at the scalp surface and retain valuable temporal information, EEG is unable to resolve the location of the source of any measured activity. This “inverse problem” is primarily driven by the heterogeneous layers of tissue, skull, and scalp with varying conductivities between the brain and scalp (i.e., between the source and measurement device). The electrical potentials measured at the scalp surface are in turn, distorted across the scalp, and without the ability to accurately model the conductivities of different layers of tissue, it is difficult to localize the measured signals across the brain. MEG is described as the magnetic equivalent to EEG due to the specific properties of magnetic fields. MEG is impacted by the inverse problem to a lesser degree compared to EEG as the measured magnetic field is not distorted by heterogeneous layers of tissue between the brain and the measuring device (Hämäläinen 1992). Source localization is considerably easier with MEG compared to EEG and there exist many possible “solutions” to the inverse problem via equivalent dipole fitting, beamforming, and Bayesian methods. However, the shortcoming of MEG is that it is only sensitive to currents tangential to the scalp surface as only these currents produce magnetic fields detectable outside the head (Hämäläinen 1992; Singh 2014). Further, activity from deep or subcortical sources is challenging to resolve with MEG. In whole, this distinction makes determining sources from MEG recordings preferred over EEG, though the spatial resolution of both of these techniques is limited compared to fMRI.

Because fMRI and MEG visualize different neural sources and measure different aspects of brain function, these methods are complementary (Wang and Holland 2014; Wang, Holland, and Vannest 2012). FMRI measures metabolic processes that drive neural firing and is able to identify primary brain regions and networks involved with different tasks with millimeter accuracy, but its poor temporal resolution limits its use to determine directional information flow within a network. MEG directly reflects magnetic fields generated by postsynaptic neural activity and preserves the millisecond time-scale of neurophysiologic activity, but unlike hemodynamic methods, it has limited spatial resolution. Implementing a multi-modal approach by combining hemodynamic and electrophysiological functional connectivity methods capitalizes on the advantages of both modalities. We are able to develop complete models of verbal memory in healthy populations by characterizing neuroanatomical contributions alongside sequential and dynamic aspects of verbal encoding with high spatiotemporal resolution. The ability to capture and describe the differential recruitment of contributing brain areas during a task is especially relevant during working memory, as the time scale of brain activity during verbal memory is known to be fast (Marinković 2004).

Successful studies combining the strengths of both MEG and fMRI have mainly focused on sensory or motor processing (Ahlfors et al. 1999; Auranen et al. 2009; Schulz et al. 2004; Stippich et al. 1998; Tuunanen et al. 2003) and rest (Lottman et al. 2019). Studies investigating higher cognitive functions are limited (Wang et al. 2012). It is especially challenging for MEG to resolve source localization that involves complex and distributed neural processes: inducted activity is not likely to be time-locked to stimuli across subjects, may arise from multiple generators across the cortex, and thus can be difficult to isolate (Lopes da Silva 2013; Takeda et al. 2014). When examining distributed responses during a cognitive task, certain considerations should be discussed. Experimental paradigms evaluating sensory responses, including visual, auditory, motor, and somatosensory responses, typically involve very short paradigms (and, in turn, have high signal-to-noise ratio (SNR) due to averaging hundreds of trials). However, it is substantially more difficult to reach similar SNR among paradigms associated with high-order cognitive tasks due to the duration of each trial (Wang and Holland 2021). Additionally, averaging across trials when investigating induced activity has the potential to weaken SNR when responses are not time-locked to the stimulus across runs, unlike consistently identified event-related potentials (ERPs) of sensory stimuli and related evoked activity (Hillebrand et al. 2005; Takeda et al. 2014).

The main goal of our study was to develop a processing stream that combines information from fMRI and MEG during a verbal memory learning and attention task to maximize the strengths of each modality. FMRI’s spatial resolution and MEG’s temporal resolution may be resolved together within a Bayesian framework by applying the Multiple Sparse Prior (MSP) algorithm (Friston et al. 2008a; Henson et al. 2011, 2019). We used this pipeline to identify spatiotemporal characteristics of active encoding with improved spatial and temporal resolution when compared to fMRI or MEG approaches alone.

## 2. METHODS

### 2.1 Participants

We recruited 24 healthy native English-speaking adults (13 female, 4 atypically handed, ages 18-39 years) with no history of neurological or psychiatric disorders (Table 1). Participation included attending two separate sessions of scanning approximately one week apart. Of the 24 participants, 23 completed all study procedures and are included in the analyses. The Institutional Review Board at the University of Alabama at Birmingham approved this project, and all participants provided written informed consent. Participants completed the MRI scanning session during their first visit and the MEG scanning session during their second visit (Nair et al. 2019; Vannest et al. 2015).

**Table 1:**
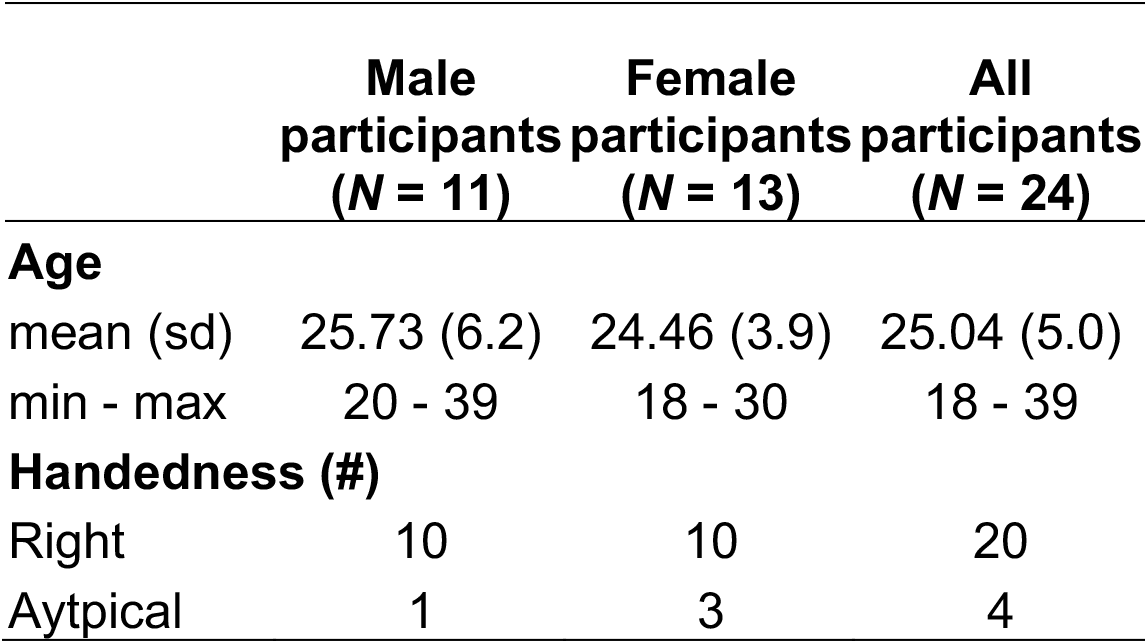
Participant demographics.

### 2.2 Paired-Associate Learning Task

During both fMRI and MEG sessions, related word pairs were presented to participants during the verbal paired-associate learning task (Schefft et al. 2008). All word pairs selected for the present study were all under 6 letters in length and were chosen from previous studies (Schefft et al. 2008; Siegel et al. 2012). Word pairs were distributed across 5 relationship classes: associates (e.g., *lock – key*), category members (e.g., *saucer – bowl*), synonyms (e.g., *street – road*), antonyms (e.g., *hot – cold*), and rhymes (e.g., *care – dare*) (Siegel et al. 2012). To ensure comprehension regarding task procedures, participants practiced a version of the task before their first scanning session. During fMRI scanning, 60 word pairs were presented either in full (e.g., *spider – web*), or with the second word partially missing (e.g., *bed – p\*\*\*\*\**), and participants were instructed to either passively read or actively self-generate the second word out loud. During MEG scanning, 200 word pairs were presented, but only with the second word partially missing and participants were instructed to *covertly* generate the target word for each trial. There was no overlap in the word pairs used in the fMRI and MEG versions of the task. For all but three participants, fMRI scanning sessions preceded MEG scanning sessions by several weeks.

### 2.3 Data Acquisition

#### 2.3.1 MRI Acquisition

Anatomical and fMRI data were acquired on a Siemens Magnetom Prisma 3.0T whole-body MRI system during approximately one hour-long scanning session. High-resolution T1-weighted anatomical images were acquired (TR: 2400ms, TE: 2.22ms, FOV: 25.6×24.0×16.7cm, matrix 256×240, flip angle: 8 degrees, slice thickness: 0.8mm, voxel size: 0.8×0.8×0.8mm). During the in-scanner pairs encoding task, functional T2*-weighted images were obtained using a partially silent event-related task design (TR: 1990ms, TE: 35ms, FOV: 240×240×129mm, flip angle: 70 degrees, matrix 240×240, slice thickness: 4mm, axial slices, voxel size: 3.8×3.8×4mm). This clustered-sparse temporal acquisition technique (HUSH; Schmithorst and Holland 2004) allows for recording overt responses during scanning while taking advantage of the intrinsic delayed response of the HRF. The HUSH technique captures activity taking place seconds preceding data collection, as the positive peak of the HRF occurs around 4 to 6 seconds following stimulus presentation (Buxton et al., 2004).

#### 2.3.2 MEG Acquisition

MEG recordings (sampling frequency of 291 Hz) were collected in a magnetically shielded room using a 148-channel whole head magnetometer (MagnesTM 2500 WH, 4-D Neuroimaging). Fiducial head-sensor coils and head shape data using a 3D digitizer were collected to monitor head position and for co-registration between participant’s MEG data and anatomical MRI. Collection of task and a resting-state scans was counterbalanced.

### 2.4 FMRI Data Processing

We have outlined the co-processing pipeline for clarity in Figure 1, and have also included in the appendix annotated MATLAB scripts (.mat) for the entire pipeline (directory creation, fMRI and MEG preprocessing and second level analyses). These can be adapted to suit various task designs. Preprocessing of event-related fMRI data was performed with SPM12 (http://www.fil.ion.ucl.ac.uk/spm/). To account for any signal intensity changes in the HRF across three volumes acquired via sparse acquisition, functional image volumes were split into three separate HUSH parts (Schmithorst and Holland, 2004), realigned and coregistered to individual participant’s anatomical before normalization to the MNI152 template atlas. All functional image volumes were then spatially smoothed to an effective smoothness of a Gaussian FWHM of 6mm. There were no additional filtering or artifact regression steps prior to Group ICA (Calhoun et al., 2001). All trials were retained for analysis as participants attempted to reach semantic and contextual integration during the task, and thus are assumed to undergo the process of encoding word pair associates regardless of whether they produce the correct word during the task (Marinkovik, 2004). Further details regarding fMRI preprocessing of HUSH data, including a broad schematic of the developed analysis pipeline, can be found in our previously published work (Nair et al. 2019).

**Figure 1:**
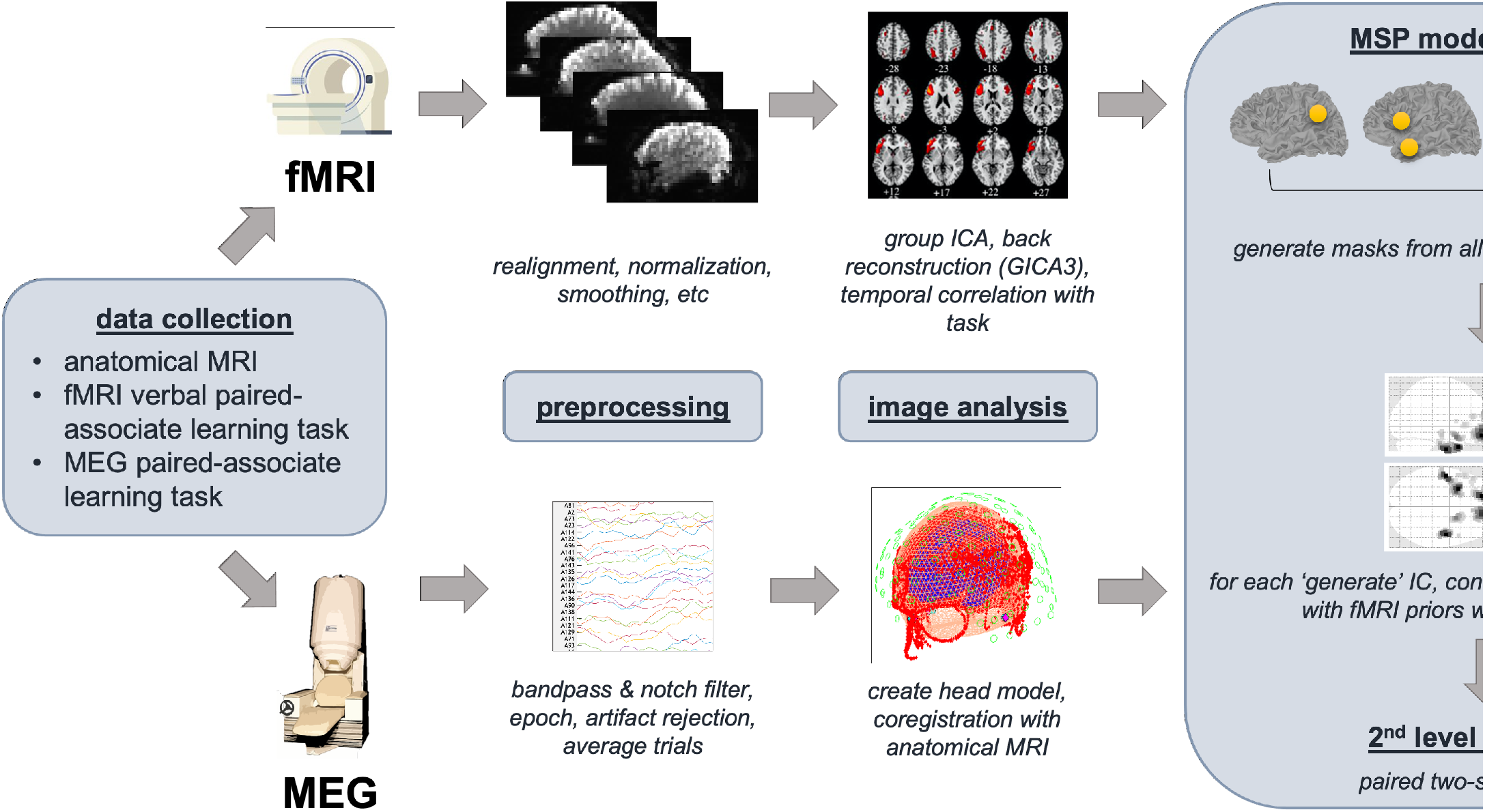
Co-processing pipeline overview/schematic. Data collection (section 2.3), preprocessing fMRI (2.4.1) and MEG (2.5.1), group level fMRI analysis using group ICA (2.4.2) and MEG image analysis (2.5.2), MSP model inversion steps (2.4.2 and 2.5.2), and second level analysis comparing “generate” and “baseline” conditions (2.5.3).

Group-level spatial independent component analysis (ICA) was carried out using Group ICA fMRI Toolbox, v4.0b (GIFT; http://mialab.mrn.org/software/gift) implemented in MATLAB (Calhoun et al. 2001). Functional image volumes were not subject to any additional filtering or artifact regression prior to Group ICA (Calhoun et al. 2001). First, two rounds of subject-specific principle component analysis (PCAs) were conducted for data reduction for each of the three HUSH image volumes. The first round of PCA at the individual subject level yielded 51 components, and the second yielded 41 components (Calhoun et al. 2001; Erhardt et al. 2011). A high model order of 41 independent components was selected using Infomax algorithm to aid in segmenting task-related brain activity into functionally distinct and noise-related sources (Hutchison and Morton 2015; Ray et al. 2013; Saliasi et al. 2014; Ystad et al. 2010) consistent with similar studies from our group (Nair et al. 2019; Vannest et al. 2015). Using GIFT’s GICA3 back-reconstruction method, subject-specific time courses and spatial maps (SMs) were estimated from matrices derived in previous PCA steps (Calhoun et al. 2001; Erhardt et al. 2011). GIFT yields these spatial maps that can be interpreted as networks of similar BOLD activity, related to various aspects of processing (Calhoun et al., 2001; McKeown et al., 2003). We used GIFT’s temporal sorting tool to classify components by comparing the model’s time course to the time courses of all 41 ICs using a correlation function (Rachakonda et al. 2007). The model time course was inputted as a binary task time series, where a “1” was used to designate the active encoding condition (task-positive trials), and a “0” identified passive reading (task-negative) trials. Components with a correlation coefficient of |r|>0.30 were selected as task-related (Gaston et al. 2017; Nair et al. 2019; Vannest et al. 2015). After temporal sorting, components are arranged based on correlation with the binary time series, and those with positive and negative correlations that meet threshold are visually examined. Of the 41 components identified, 4 components met threshold for task-positive relatedness, and 1 component was identified as task-negative. For the active encoding condition, component correlations were *r*>0.30 and for the passive reading condition, correlations were *r*>-0.30. Components with a correlation coefficient between -0.30>*r*>0.30 were excluded from any further analyses.

To resolve any discrepancies using the HUSH sparse acquisition paradigm and three GIFT analyses, task-related components were visually inspected to identify similar regional activity across HUSH parts. When a component met task-relatedness threshold across more than one volume, then the volume with the highest correlation to the task time series was selected. Once components were selected as meeting threshold for both task-positive and -negative conditions, masks were directly created from group ICA NIFTI files using AFNI’s ‘Save Mask’ tool in the graphical user interface.

### 2.5 MEG Data Processing

MEG raw data in 4D/BTI format were first converted using SPM’s convert function (spm_eeg_convert), and headshape points were incorporated (spm_eeg_prep). An ERP approach was employed in order to identify early sensory responses to active generation, and an approach isolating theta frequency band (4-7Hz) was employed due to the lexical-semantic retrieval nature of the task (Bastiaansen et al. 2005; Klimesch 1999; Pu et al. 2020; Raghavachari et al. 2006).

#### ERP approach

A series of filters was applied to the converted raw data (spm_eeg_filter): a high pass filter at 0.6Hz, followed by a notch filter to remove line noise (60Hz, 120Hz), and a low pass filter at 30Hz. Data were epoched (spm_eeg_epochs), trials were averaged (spm_eeg_average), and a low pass filter at 30Hz was applied once more due to the potential introduction of high frequencies after averaging (Litvak et al. 2011).

#### Theta power approach

To analyze event-related theta power, additional analyses averaging power across the time-frequency window (4-7Hz) were conducted (Henson et al. 2019). For these data, a high pass filter at 0.6Hz, notch filters at 60Hz and 120Hz, and a low pass filter at 60Hz were applied. To preserve cognitive activity that is not likely time-locked across trials, within subject epochs were not averaged before moving forward with source localization.

For both approaches, artefact detection was conducted using SPM’s threshold z-scored data detection algorithm with the threshold set at z=3 (spm_eeg_artefact). Artefacts were marked as events with an excision window around each event of 100ms. Channels with more than 80% of trials with detected artefacts were declared as bad. Data were epoched using a trial definition file that establishes a condition label, event type, event value, and trial shift.

### 2.6 Multiple Sparse Priors: fMRI Constrained MEG

Combining data from two modalities with different neural sources is challenging. At present, there is no optimal solution for integrating MEG and fMRI data. We employed a hierarchical Bayesian approach, Multiple Sparse Prior (MSP) to approximate current density and determine possible source solutions for a verbal memory task (Friston et al. 2008a; Henson et al. 2010, 2011, 2019; Litvak et al. 2011). We used the fMRI task-positive independent components outlined in Section 2.4 as spatial priors to constrain MEG source estimations within MSP.

The MEG imaging pipeline for source analysis is divided into four steps, three of which are executed in the “g_source_reconstruction.m” script. First, source space modeling created a head model based on an individual’s structural image and corresponding spatial deformation field. A nonlinear transformation was applied to create individual cortical meshes in the designated template space using a “normal” cortical mesh size (5124 vertices). Next, data co-registration utilized the fiducials by projecting the MEG data onto each participant’s anatomical MRI using a rigid-body transformation (Henson et al. 2010). When a subject’s structural MRI is available, it is recommended to specify fiducial points but not use the head shape information (Litvak et al. 2011). Forward volume head modeling of the projected magnetic field was determined by using a single-shell (surface) fit to the individual’s scalp mesh. Lead field matrices calculated during this stage are later used for inversions. Following source space modeling and co-registration and forward computation variables are created for each fMRI prior (four task-positive IC’s). Source reconstruction takes place during the model inversion stage of processing, using multiple sparse priors for each prior using greedy search (GS) inversion type, with a full time-frequency window of [-100 to 1200 ms] and a prior mask (.nii) for each prior derived fMRI components from section 2.4.3.

In the final stage of source analysis, inverse reconstruction, time-frequency contrasts were created by inverting the epoch of interest (across listed frequencies) to localize the effects within the cortical mesh (Henson et al. 2019). For the ERP approach, time windows analyzed (in ms) included [0 150], [100 250], [300 450], [350 500], [500 650], [600 750], and [700 850], and a baseline condition at [-150 0]. For the theta approach, time windows included [100 250], [300 450], [450 600], [600 750], [750 900], [900 1050], [1050 1200] with frequencies restricted to 4-7 Hz.

To examine spatial projections at the group level, paired two-sample t-tests were conducted for each prior and for each time window of interest, contrasting the active generation condition with a baseline condition (stats.factorial_design).

## 3. RESULTS

### 3.1 Group ICA: Task-related components from fMRI data

Five components met threshold for task-relatedness out of the 41 components identified using Group ICA (Table 2, Figure 2). Four were the active generation condition (positive correlation with task time course) and one was the passive reading condition (negative correlation with task time course). The component with the highest task correlation (IC19; *r*=0.4142) included regions spanning right supramarginal gyrus, postcentral gyrus, and middle frontal gyrus, bilateral superior and inferior parietal lobes (IPL), and precuneus (Figure 2a). The component with the second highest task-positive correlation (IC13; *r*=0.3674) included regions spanning bilateral inferior, medial, and middle frontal gyri, supramarginal gyri, inferior and superior parietal lobes, left superior temporal gyrus (STG) and middle temporal gyrus (MTG), superior frontal gyrus, precentral gyrus, and precuneus (Figure 2b). The third component (IC4; *r*=0.3652) included bilateral cuneus, inferior and middle occipital gyri, lingual gyri, and supramarginal gyri (Figure 2c). The fourth task-positive component (IC26; *r*=0.3319) spans bilateral inferior and superior frontal gyri, insula, anterior cingulate cortex (ACC) and medial frontal gyri, STG, and extra-nuclear areas, and right middle frontal gyrus and caudate (Figure 2d). The one component meeting task-relatedness for passive reading (IC38; *r*=-0.3466) spans bilateral precuneus, cingulate and posterior cingulate cortices (PCC), angular gyri, STG and MTG, inferior and superior parietal lobe, and left superior and middle frontal gyri, and right supramarginal gyrus and extra-nuclear areas (Figure 2e). All four task-positive components were retained as fMRI priors (binarized masks) for MEG source reconstruction, and the distributed cortical solutions for each prior’s model inversion are listed below.

**Table 2:**
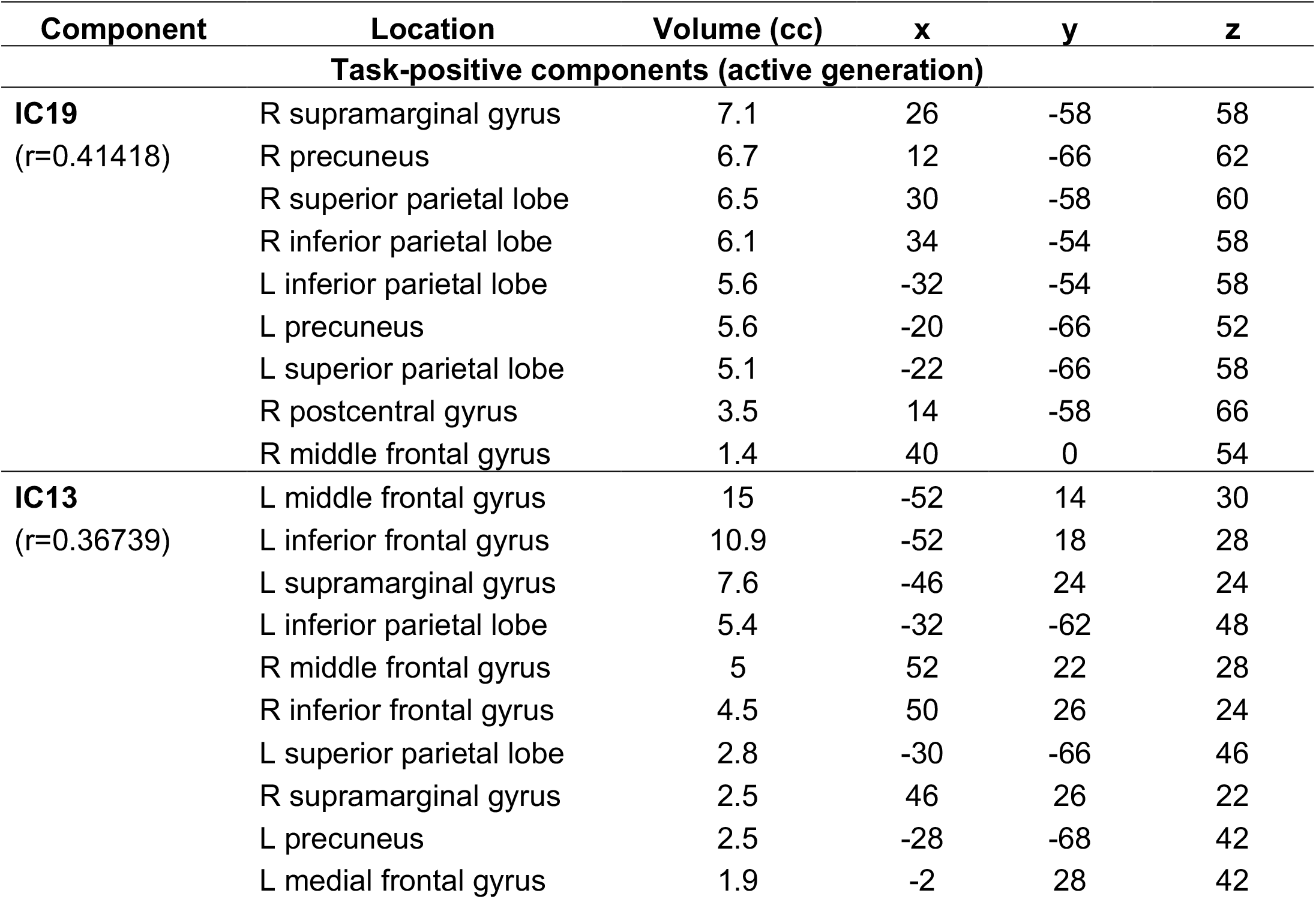
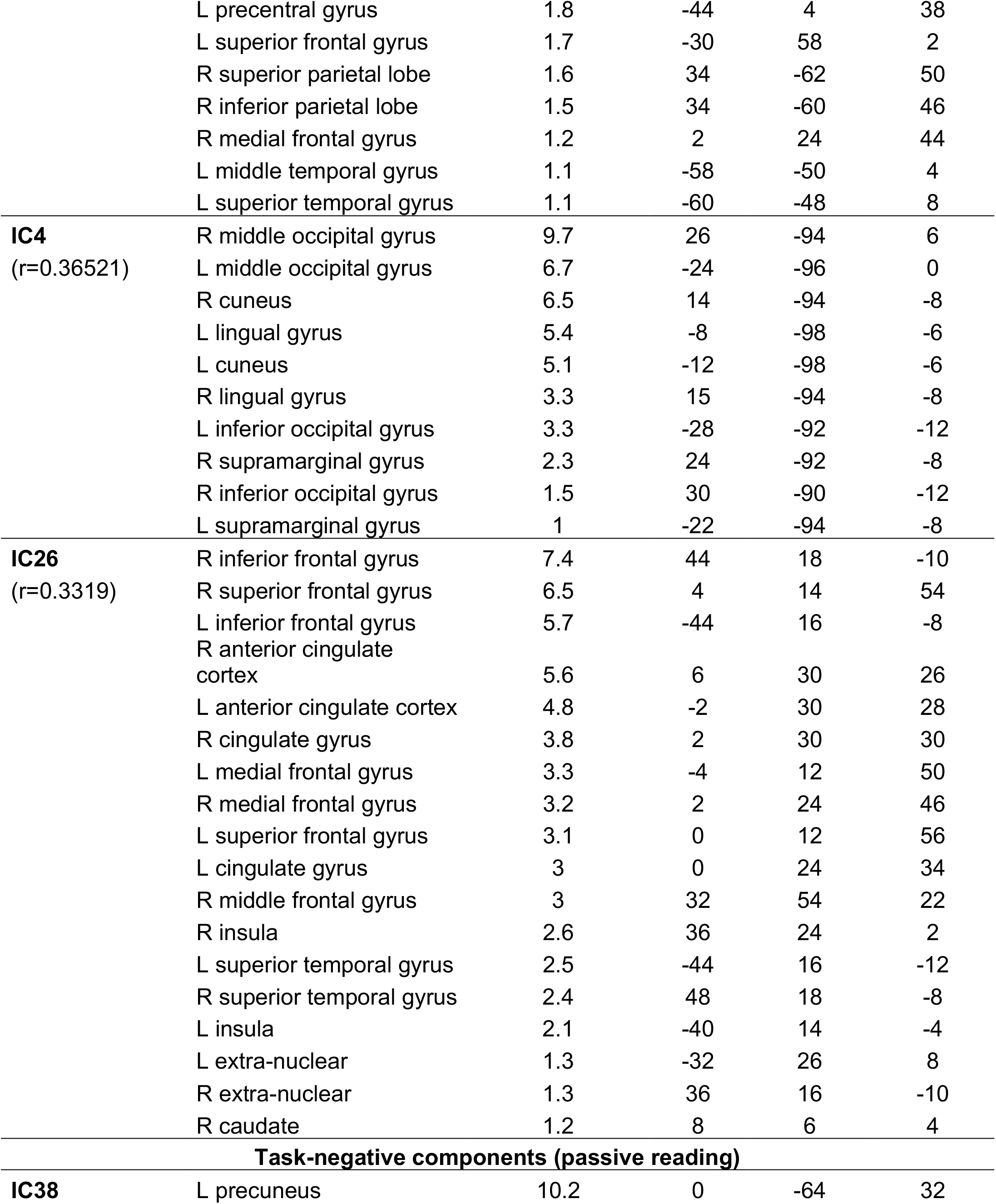
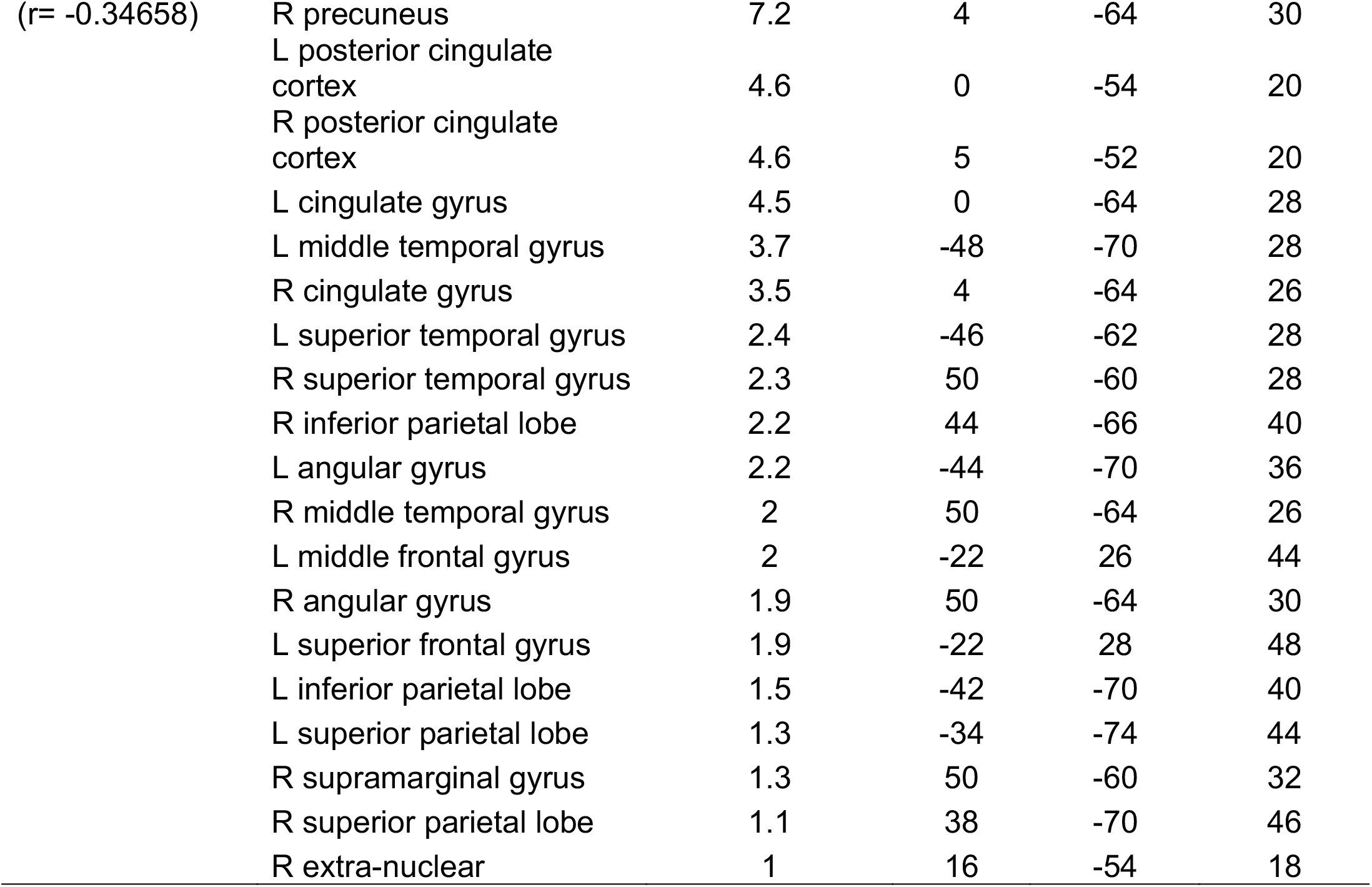
Group level task-related ICs from fMRI (Figure 2) and the location, volume and MNI coordinates (x, y, z) for each IC cluster’s center of mass. L, left; R, right.

| Component | Location | Volume (cc) | x | y | z |
| --- | --- | --- | --- | --- | --- |
| **Task-positive components (active generation)** |  |  |  |  |  |
| **IC19**<br>(*r*=0.41418) | R supramarginal gyrus | 7.1 | 26 | -58 | 58 |
|  | R precuneus | 6.7 | 12 | -66 | 62 |
|  | R superior parietal lobe | 6.5 | 30 | -58 | 60 |
|  | R inferior parietal lobe | 6.1 | 34 | -54 | 58 |
|  | L inferior parietal lobe | 5.6 | -32 | -54 | 58 |
|  | L precuneus | 5.6 | -20 | -66 | 52 |
|  | L superior parietal lobe | 5.1 | -22 | -66 | 58 |
|  | R postcentral gyrus | 3.5 | 14 | -58 | 66 |
|  | R middle frontal gyrus | 1.4 | 40 | 0 | 54 |
| **IC13**<br>(*r*=0.36739) | L middle frontal gyrus | 15 | -52 | 14 | 30 |
|  | L inferior frontal gyrus | 10.9 | -52 | 18 | 28 |
|  | L supramarginal gyrus | 7.6 | -46 | 24 | 24 |
|  | L inferior parietal lobe | 5.4 | -32 | -62 | 48 |
|  | R middle frontal gyrus | 5 | 52 | 22 | 28 |
|  | R inferior frontal gyrus | 4.5 | 50 | 26 | 24 |
|  | L superior parietal lobe | 2.8 | -30 | -66 | 46 |
|  | R supramarginal gyrus | 2.5 | 46 | 26 | 22 |
|  | L precuneus | 2.5 | -28 | -68 | 42 |
|  | L medial frontal gyrus | 1.9 | -2 | 28 | 42 |
|  | L precentral gyrus | 1.8 | -44 | 4 | 38 |
|  | L superior frontal gyrus | 1.7 | -30 | 58 | 2 |
|  | R superior parietal lobe | 1.6 | 34 | -62 | 50 |
|  | R inferior parietal lobe | 1.5 | 34 | -60 | 46 |
|  | R medial frontal gyrus | 1.2 | 2 | 24 | 44 |
|  | L middle temporal gyrus | 1.1 | -58 | -50 | 4 |
|  | L superior temporal gyrus | 1.1 | -60 | -48 | 8 |
| **IC4**<br>(r=0.36521) | R middle occipital gyrus | 9.7 | 26 | -94 | 6 |
|  | L middle occipital gyrus | 6.7 | -24 | -96 | 0 |
|  | R cuneus | 6.5 | 14 | -94 | -8 |
|  | L lingual gyrus | 5.4 | -8 | -98 | -6 |
|  | L cuneus | 5.1 | -12 | -98 | -6 |
|  | R lingual gyrus | 3.3 | 15 | -94 | -8 |
|  | L inferior occipital gyrus | 3.3 | -28 | -92 | -12 |
|  | R supramarginal gyrus | 2.3 | 24 | -92 | -8 |
|  | R inferior occipital gyrus | 1.5 | 30 | -90 | -12 |
|  | L supramarginal gyrus | 1 | -22 | -94 | -8 |
| **IC26**<br>(r=0.3319) | R inferior frontal gyrus | 7.4 | 44 | 18 | -10 |
|  | R superior frontal gyrus | 6.5 | 4 | 14 | 54 |
|  | L inferior frontal gyrus | 5.7 | -44 | 16 | -8 |
|  | R anterior cingulate cortex | 5.6 | 6 | 30 | 26 |
|  | L anterior cingulate cortex | 4.8 | -2 | 30 | 28 |
|  | R cingulate gyrus | 3.8 | 2 | 30 | 30 |
|  | L medial frontal gyrus | 3.3 | -4 | 12 | 50 |
|  | R medial frontal gyrus | 3.2 | 2 | 24 | 46 |
|  | L superior frontal gyrus | 3.1 | 0 | 12 | 56 |
|  | L cingulate gyrus | 3 | 0 | 24 | 34 |
|  | R middle frontal gyrus | 3 | 32 | 54 | 22 |
|  | R insula | 2.6 | 36 | 24 | 2 |
|  | L superior temporal gyrus | 2.5 | -44 | 16 | -12 |
|  | R superior temporal gyrus | 2.4 | 48 | 18 | -8 |
|  | L insula | 2.1 | -40 | 14 | -4 |
|  | L extra-nuclear | 1.3 | -32 | 26 | 8 |
|  | R extra-nuclear | 1.3 | 36 | 16 | -10 |
|  | R caudate | 1.2 | 8 | 6 | 4 |
| **Task-negative components (passive reading)** |  |  |  |  |  |
| **IC38** | L precuneus | 10.2 | 0 | -64 | 32 |
| (r= -0.34658) | R precuneus | 7.2 | 4 | -64 | 30 |
|  | L posterior cingulate cortex | 4.6 | 0 | -54 | 20 |
|  | R posterior cingulate cortex | 4.6 | 5 | -52 | 20 |
|  | L cingulate gyrus | 4.5 | 0 | -64 | 28 |
|  | L middle temporal gyrus | 3.7 | -48 | -70 | 28 |
|  | R cingulate gyrus | 3.5 | 4 | -64 | 26 |
|  | L superior temporal gyrus | 2.4 | -46 | -62 | 28 |
|  | R superior temporal gyrus | 2.3 | 50 | -60 | 28 |
|  | R inferior parietal lobe | 2.2 | 44 | -66 | 40 |
|  | L angular gyrus | 2.2 | -44 | -70 | 36 |
|  | R middle temporal gyrus | 2 | 50 | -64 | 26 |
|  | L middle frontal gyrus | 2 | -22 | 26 | 44 |
|  | R angular gyrus | 1.9 | 50 | -64 | 30 |
|  | L superior frontal gyrus | 1.9 | -22 | 28 | 48 |
|  | L inferior parietal lobe | 1.5 | -42 | -70 | 40 |
|  | L superior parietal lobe | 1.3 | -34 | -74 | 44 |
|  | R supramarginal gyrus | 1.3 | 50 | -60 | 32 |
|  | R superior parietal lobe | 1.1 | 38 | -70 | 46 |
|  | R extra-nuclear | 1 | 16 | -54 | 18 |

**Figure 2:**
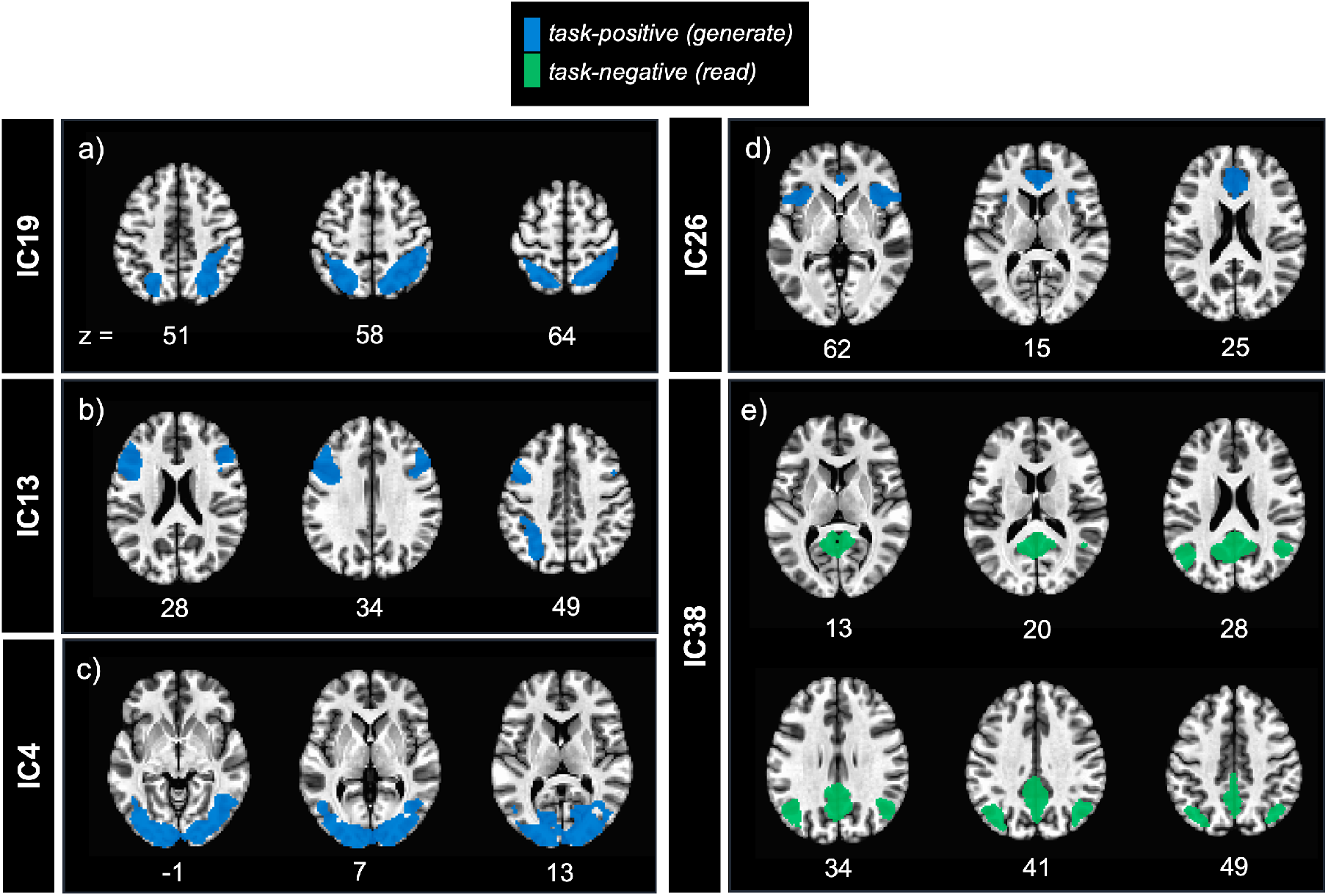
Task-related fMRI components. Images are presented in neurological orientation (right in the image is right in the brain). (A-F) are group average independent components positively correlated (*r*>0.25) and (G) is negatively correlated (*r*<0.25) with the time course. Components span: (A) bilateral SPL and IPL, precuneus, middle occipital gyri, and right supramarginal gyrus (IC19); (B) bilateral IFG, precentral gyri, left superior frontal gyrus, middle orbital gyrus, IPL, middle occipital gyrus (IC13); (C) bilateral calcarine gyri, fusiform gyri, lingual gyri, middle occipital gyri, inferior temporal gyri (IC4); (D) bilateral IFG, insula, ACC, MCC, supplementary motor area (IC26); (E) bilateral precuneus, MCC, calcarine and angular gyri, PCC, middle occipital gyri, MTG (IC38). SPL: superior parietal lobe; IPL: inferior parietal lobe; IFG: inferior frontal gyrus; ACC: anterior cingulate cortex; MCC: middle cingulate cortex; MTG: middle temporal gyrus; PCC: posterior cingulate cortex

### 3.2 MEG Source Reconstruction

Source reconstruction was performed in standardized MNI space for each fMRI prior resulting in summarized images (Figures 3 and 4). Reconstructions were run for all four fMRI priors (independent components meeting task-positive threshold) for each subject, producing individual subject spatial maps. Results from the paired two-sample t-test contrasting the active generation condition to baseline period are presented in Figures 3b-c, 4b-c. Source reconstruction maps were also generated without fMRI priors for the ERP and theta power (Figures 3a, 4a) approaches. All results are corrected for multiple comparisons at *p*<0.05.

**Figure 3:**
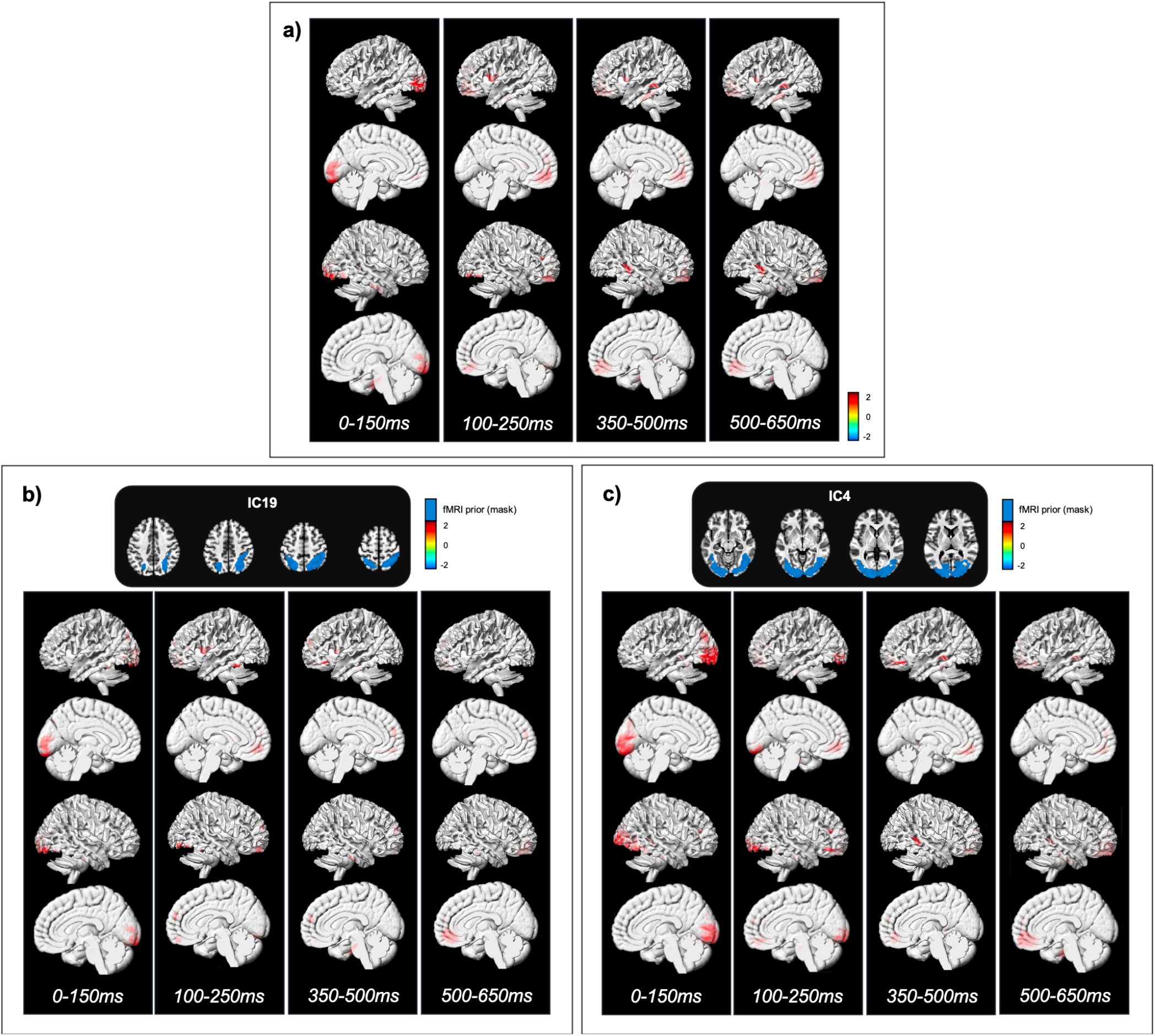
Multiple sparse priors (MSP) for ERP approach. (a) no priors approach, (b) IC19 constrained source reconstructions, (c) IC4 constrained reconstructions. Images are presented in neurological orientation (R=R).

**Figure 4:**
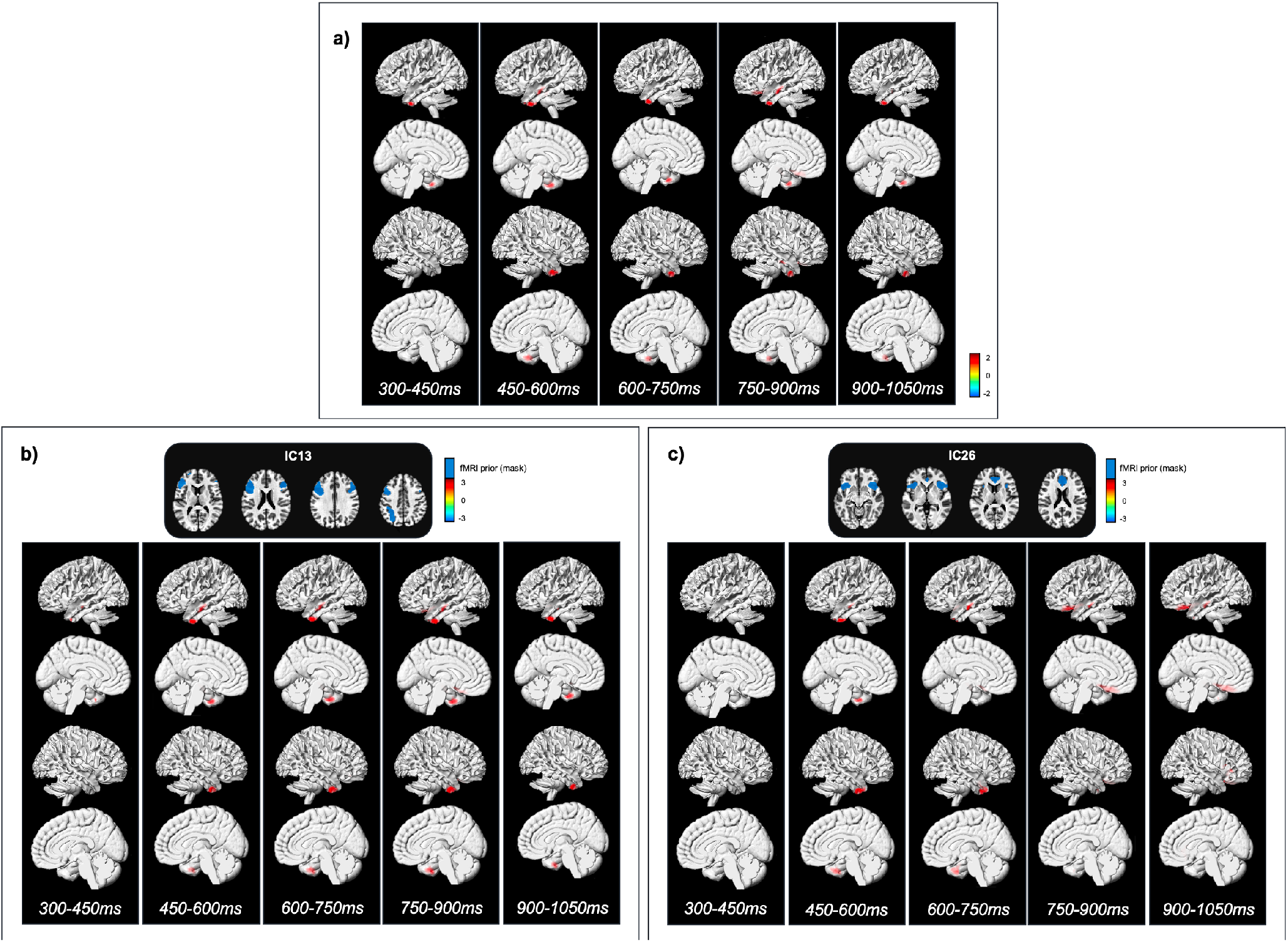
Multiple sparse priors (MSP) for event-related theta-power approach. (a) no priors approach, (b) IC13 constrained source reconstructions, (c) IC26 constrained reconstructions. Images are presented in neurological orientation (R=R).

#### IC19 (Figure 3b)

Source reconstruction results using the ERP approach for IC19 using fMRI prior that spans bilateral parietal areas are presented in Figure 3b and Table 3. Early activity in the first 150ms across bilateral occipital and left inferior temporal gyri, followed by recruitment of bilateral inferior temporal gyri, superior orbitofrontal and medial frontal gyri between 100 and 250ms. Between 350 and 500ms, group activity spans medial and inferior frontal gyri, right parahippocampal gyrus, and left inferior temporal gyrus, and between 500 and 650ms, sustained activity in bilateral medial frontal gyrus right medial orbitofrontal cortex (OFC), and fusiform gyrus.

**Table 3:**
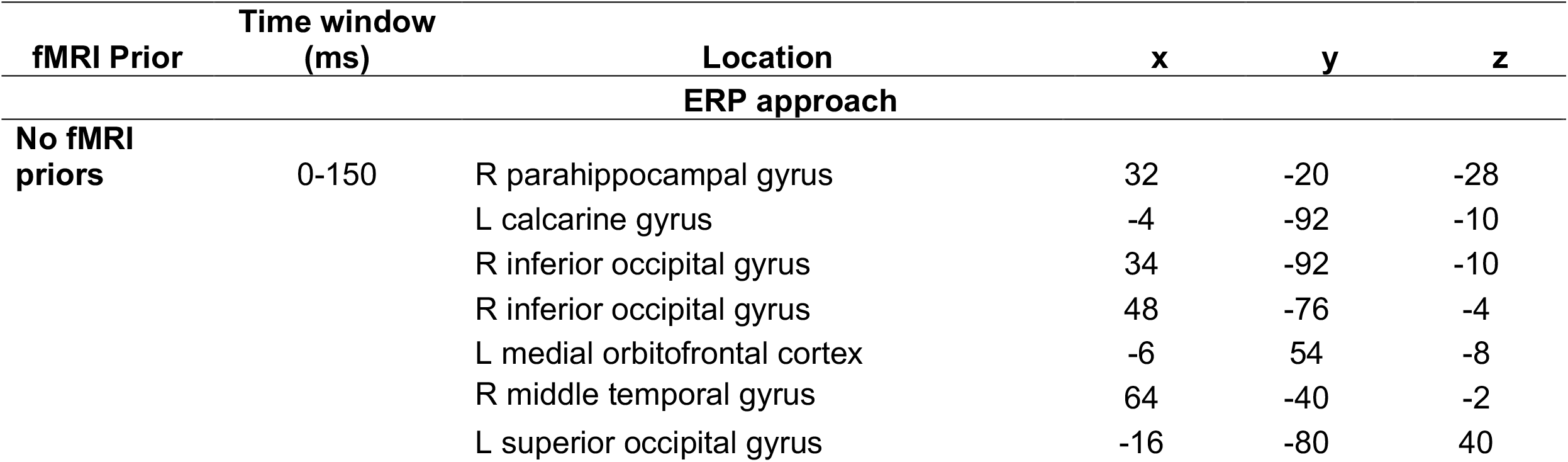
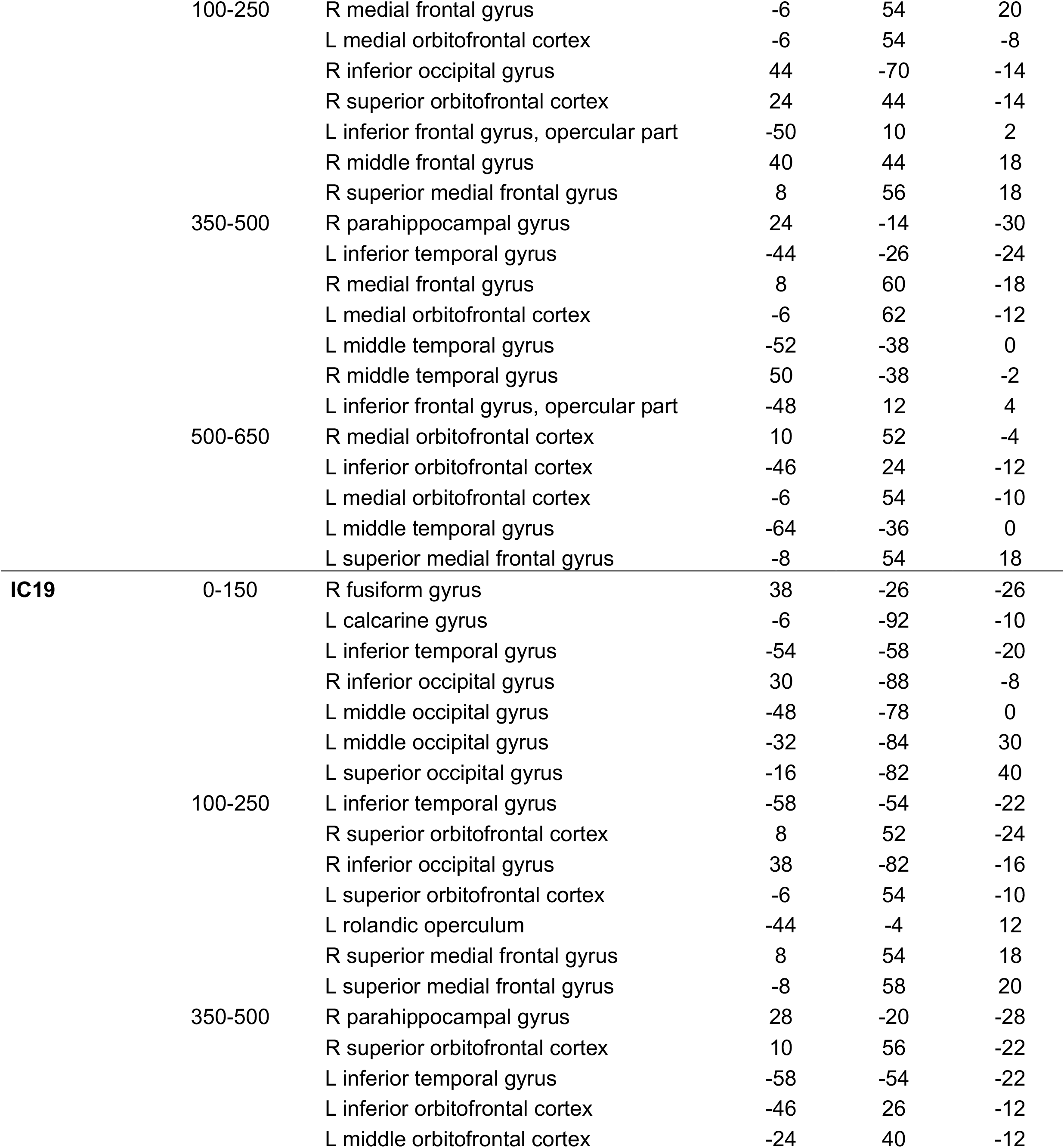
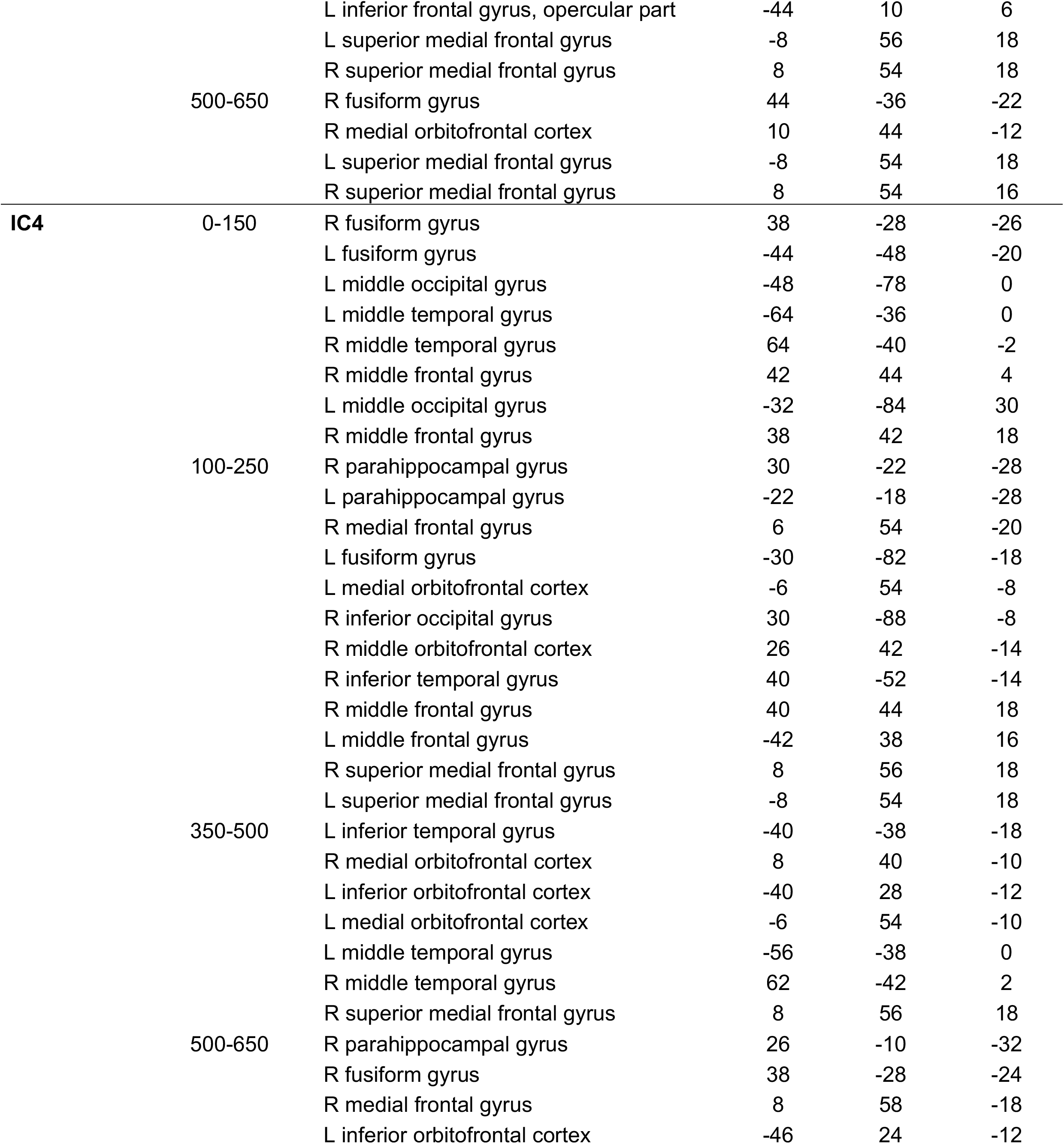
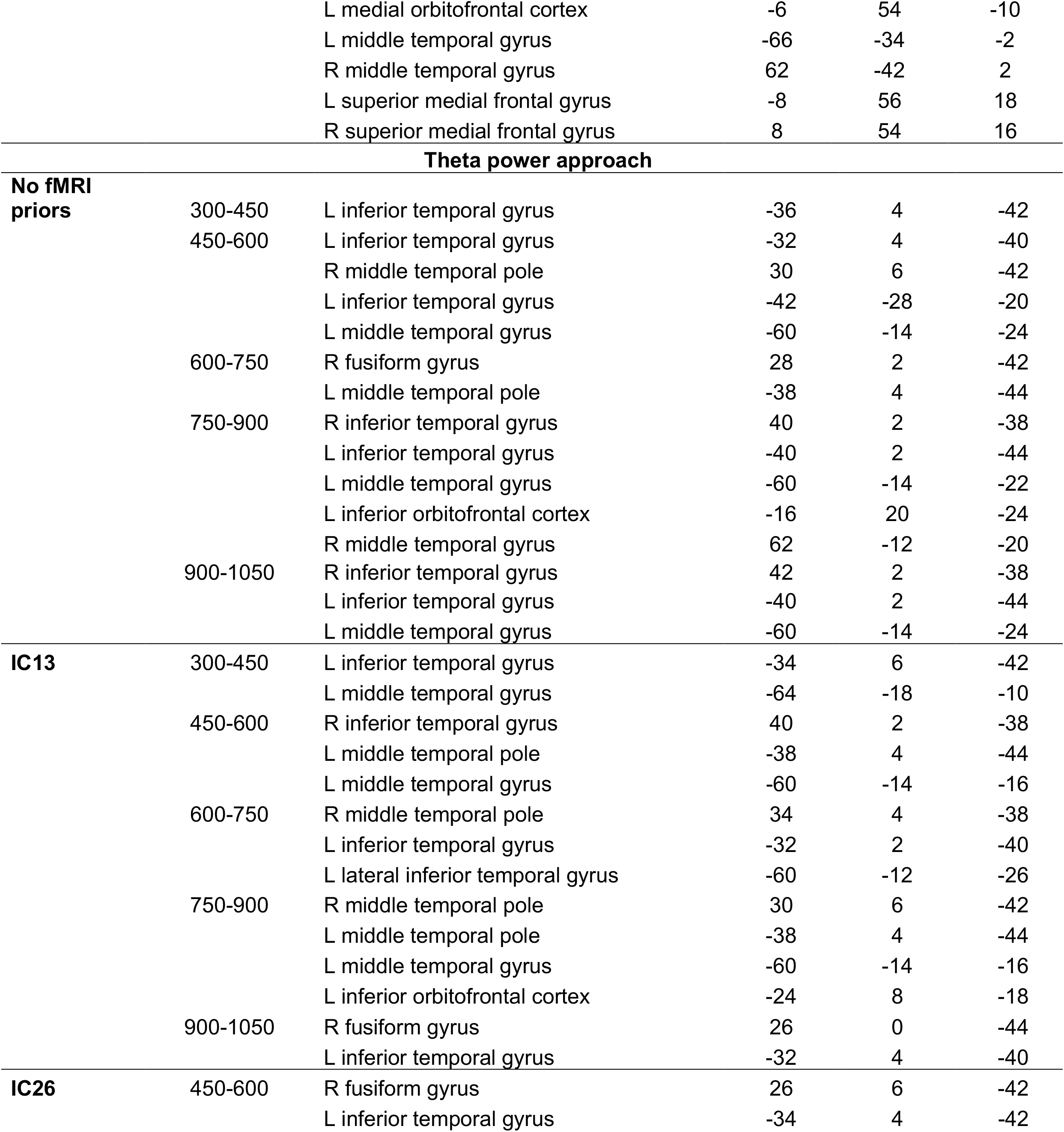
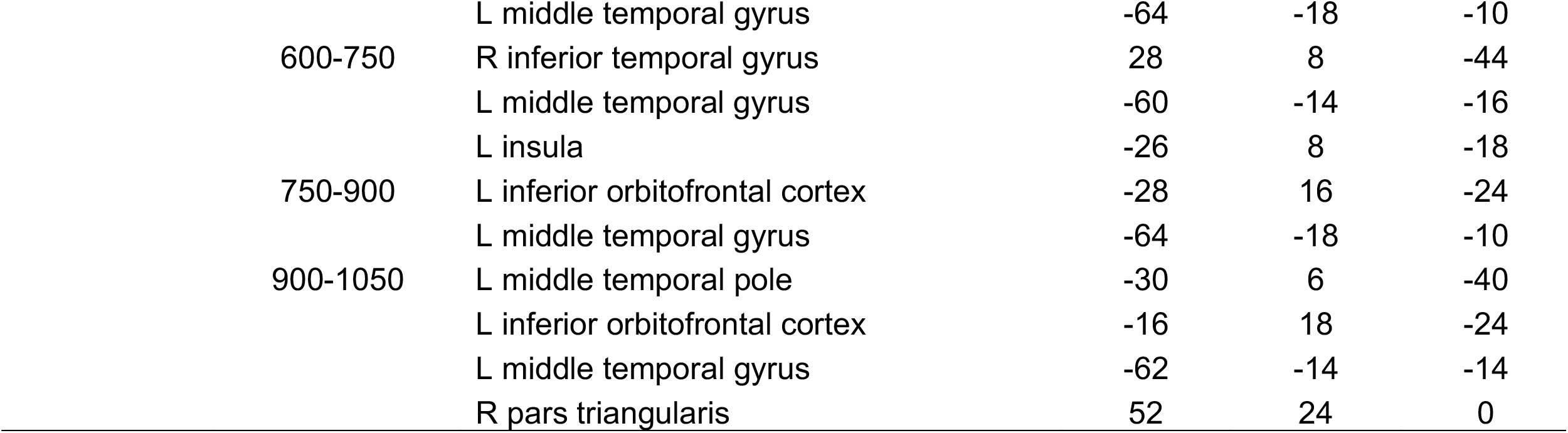
MEG source reconstruction results for each fMRI prior using the ERP approach (Figure 3) and the theta-power approach (Figure 4). For each time window, the location and MNI coordinates (x, y, z) for each source location’s center of mass are provided. L, left; R, right.

#### IC13 (Figure 4b)

Source reconstruction images for our theta power approach for IC13 using fMRI prior that includes bilateral frontal gyri and left parietal lobe are presented in Figure 4b and Table 3. Early activity between 300 and 450ms in left inferior temporal gyrus and MTG, followed by right inferior temporal gyri and left middle temporal pole between 450 and 600ms. Within the next 150ms, left MTG joins right middle temporal pole. The time window between 750 and 900ms yielded broad, bilateral temporal activity and left inferior OFG. From 900 to 1050ms, we see right fusiform and left inferior temporal gyri contributions.

#### IC4 (Figure 3c)

Source reconstruction results using the ERP approach for IC4 using fMRI prior that spans bilateral occipital areas are presented in Figure 3c and Table 3. Activity within bilateral fusiform gyri, MTG, right middle frontal gyri, and strong middle occipital gyrus contributions are present within the first 150ms, with bilateral parahippocampal gyrus, superior medial and middle frontal gyri and orbitofrontal gyri following between 100-250ms. Within the 350 to 500ms window, we see bilateral MTG recruited, along with left inferior and medial orbitofrontal gyri, and right superior medial frontal gyrus. Within 500 to 650ms, there is still recruitment of bilateral MTG and superior medial frontal areas, along with right fusiform gyrus.

#### IC26 (Figure 4c)

Source reconstruction images for the theta power approach for IC26 using fMRI prior within bilateral inferior frontal gyrus and anterior cingulate cortices are presented in Figure 4c and Table 3. The earliest significant activity is within the time window of 450-600ms within left inferior temporal gyrus and MTG, and right fusiform gyrus. Between 600 and 750ms, there is recruitment of right inferior temporal gyrus along with left MTG and left insula, followed by left inferior OFG and left MTG within 750 and 900ms. Activity within the left middle temporal pole, inferior OFG, and MTG, along with right pars triangularis is present within the 900 to 1050ms window.

## 4. DISCUSSION

The aim of the present study was to develop a processing pipeline that would integrate fMRI and MEG data and to use this pipeline to analyze a paired associate learning task conducted with the same group of subjects. By first establishing spatial contributions to active generation using fMRI, we are able to constrain MEG source space solutions to specific volumes of interest (Friston et al. 2008b; Henson et al. 2019).

### 4.1 fMRI: Spatial contributions to active generation

The four independent components meeting a task-positive threshold of *r*>0.30 broadly capture various aspects of active encoding. The network with the highest correlation with the task, IC19 (Figure 3b), includes bilateral inferior and superior parietal areas, which have been shown to be recruited during tasks involving maintaining visual and cognitive attention (Otten, Henson, and Rugg 2001b). The inferior parietal lobe (IPL) is thought to play a substantial role in maintaining attention (Rueckert and Grafman 1998; Rushworth, Krams, and Passingham 2001). In the present study, IPL contributions seen during self-generation (Figures 2a-b) suggest that attentional demands may be greater when actively encoding the second word in the word pair compared to passive reading.

The network with the second highest correlation with the task, IC13 (Figure 4b), is the only component meeting threshold that showed a left-lateralization effect within a broad fronto-parietal network. There is evidence suggesting the middle frontal gyri’s (MFG) role in integrating between dorsal and ventral attention streams, aiding in reorienting attention towards relevant task demands (Christensen et al. 2012; Japee et al. 2015). In the case of active generation, top-down modulation is necessary in the mental search for target words using cues, and frontal contributions likely play an integrative role during the present task. In addition to dynamic interactions with posterior brain areas, the MFG (Miller and Cohen 2001), and more broadly the dorsolateral prefrontal cortex (dlPFC), has been long affiliated with working memory processes and retrieval (D’Esposito et al. 1997; Thompson-Schill et al. 1997).

Another task-positive network with frontal contributions, IC26 (Figure 4c), shows bilateral frontal and superior temporal activity at the group level, along with ACC. This is largely consistent with previous studies of self-generation effects, where the ACC, a part of the salience network, may be playing a role in the present task when internally monitoring potential responses to the task demands (Nair et al. 2019; Rosner, Elman, and Shimamura 2013; Vannest et al. 2015). Additionally, there is recent evidence that activity within the ACC and left inferior frontal gyrus (IFG), along with superior parietal areas, may be modulated by the complexity of processing during a semantic task (Moss et al. 2011). In the present study, all three regions are implicated within two task-positive networks, suggesting increased task demand within self-generation compared to passive reading. Our findings of bilateral superior temporal gyrus involvement is also consistent with previous work on word and pseudoword recognition (Simos et al. 2000).

The third task-positive network spans bilateral visual areas, including middle and inferior occipital gyri, lingual and supramarginal gyri, and cuneus. Our findings of an extended visual network recruited during self-generation are important considering the visual nature of the task presented (Nair et al. 2019). Additionally, the differences in stimuli for passive reading and active generation may contribute to this network meeting task-relatedness as well: while the “read” condition fully presented two related word pairs on screen, the “generate” condition replaced letters in the second word pair with asterisks characters, possibly introducing variability in the visual complexity between stimuli utilized in the two conditions.

Together, brain areas associated with the default mode network (DMN) are thought to contribute to internal processing, and studies of self-generation have implicated certain areas of the DMN (IPL, medial PFC, and precuneus) to aspects of internally generating semantic information (Rosner et al. 2013). Here, we also note contributions from bilateral IPL, medial PFC, and precuneus across task-positive networks (Figures 2a-b, d) during self-generation suggesting aspects of the task encourage internal mediation perhaps during semantic and conceptual processing while searching for the correct word pair.

The only task-negative component to meet threshold for passive reading implicated a temporo-parietal network spanning PCC, superior and middle temporal gyri, inferior and superior parietal lobes. This network appears more classically aligned with the DMN, with strong activation in bilateral precuneus, IPL, and PCC, and is consistent with previous work examining brain networks associated with reading and generating (Rosner et al. 2013). In terms of the differential DMN contributions during self-generation and passive reading, it is plausible that various aspects of internal processing may recruit regionally specific areas of the DMN depending on task demands (Buckner and Carroll 2007; Shimamura 2011; Spreng, Mar, and Kim 2009).

### 4.2 MEG: ERP approach to active generation

MEG source reconstructions for the evoked approach constrained using fMRI priors (Figure 3a) and without priors (Figures 3b-c), all show early primary visual activity, followed by left inferior frontal gyrus and left inferior temporal lobe. Left IFG activity is more pronounced in the reconstruction for IC19, the fMRI network that spans bilateral parietal areas (Figure 2a) within 100-250ms. The left IFG has been long implicated in studies involving categorization of words (Demb et al. 1995; Kapur et al. 2016), and when identifying words among competing alternatives (Thompson-Schill et al. 1997; Thompson-Schill, D’Esposito, and Kan 1999). Despite parietal activity being absent from MEG reconstructions, these findings also suggest early communication within a frontoparietal network during active generation. This network has been historically associated sustained attention during a task (Ptak 2012), and has been implicated in stimulus-driven action (Corbetta, Patel, and Shulman 2008), rule-learning (de Diego Balaguer et al. 2007), and recently been found to respond to cue-related stimuli (Macaluso and Doricchi 2013).

Activity in the left OFC is more robust within 350-500ms in reconstructions for IC4 (Figure 3c), the prior spanning primary visual areas, compared to other evoked reconstructions. The timing of this activity is consistent with MEG findings of distributed, left-lateralized activity within frontopolar areas during semantic processing around 400ms (Halgren et al. 2002). These findings also support previous work suggesting prefrontal areas play a role in modulating sensory cortices during learning in a top-down fashion (Gilbert and Li 2013; Liu et al. 2020; Poort et al. 2015; Zhang et al. 2014).

### 4.3 MEG: Event-related Theta Power

Source reconstructions for the theta power approach with (Figure 4a) and without fMRI priors (Figures 4b-c) all show early left ITG activity within the first 600ms, but fMRI constrained reconstructions seemed to elucidate dynamics and information transfer more clearly than the reconstructions not informed by fMRI priors. Reconstructed time windows loosely constrained to IC26 (Figure 4c), the bilateral network including IFG, medial frontal cortex, and STG, suggest recruitment of a broad frontotemporal network during active generation. There is a clear progression of early activity from bilateral inferior temporal areas to sustained left inferior OFC and MTG activity at around 750ms, and right IFG activity after 900ms.

In relation to language processing and the generation effect, the left ITG is a critical region in the semantic system (Binder et al. 2009; Kim et al. 2011), involved in image generation (D’Esposito et al. 1997), and implicated as playing a role in stimulus encoding during visual working memory tasks (Woloszyn and Sheinberg 2009). Our findings correspond with previous work both within the temporal window and spatial characteristics of left ITG (Dhond et al. 2001; Marinkovic et al. 2012), and work suggesting left temporal theta may be modulated by retrieving lexical-semantic properties of specific words (Bastiaansen et al. 2005). The later involvement of left OFC suggests increased attention and maintenance of semantic or lexical information (Rosner et al. 2013; Tops and Boksem 2011), and along with bilateral temporal activity may represent communication underlying response selection and decision making (Young and Shapiro 2011) during active generation.

Some caution should be used when interpreting timing associated with theta-related effects, as the temporal resolution of lower frequencies is poorer than higher frequencies by roughly a few hundred milliseconds (Knösche and Bastiaansen 2002), which may contribute to the sustained activity in bilateral temporal areas across time windows (IC13, Figure 4b).

### 4.4 Differences between MSP with and without priors

Previous work examining the impact of fMRI priors on reconstructed MEG source activity has yielded mixed findings, with the MSP framework demonstrating greater model evidence compared to other techniques (Friston et al. 2008b; López et al. 2014; R.N. Henson et al. 2009), with the use of both consistent and inconsistent fMRI priors still increasing the accuracy of MEG source reconstruction (Henson et al. 2011; Wang and eHolland 2021), and priors derived from a meta-analysis also improving MEG source reconstruction (Suzuki and Yamashita 2021).

Previously, Wang and colleagues had demonstrated both spatial concordance and disagreement between MEG and fMRI among different brain areas (Wang et al. 2012). These findings suggest a model integrating the two modalities must prove robust against invalid fMRI spatial priors. MSP addresses this concern by applying a “soft” constraint of priors, which allows for disagreement between the two sources (Wang and Holland 2014). A recent study from the same group found improvement in induced activity over evoked activity during a high-order cognitive paradigm when incorporating fMRI priors into the MEG inversion solution (Wang and Holland 2021). These findings together highlight the ability of MSP to select either a sparse solution or multiple, distributed cortical sources automatically in a data driven way, using empirical priors.

In the current study, source reconstructions without fMRI priors yielded more distributed solutions (Figure 2) compared to reconstructions constrained by fMRI priors. Overall, MSP solutions for both ERP and theta-power approaches produced more focused regions that were largely, but not entirely, present in solutions without priors. This can be seen with early, sustained (0-500ms) left inferior temporal gyrus activity in solutions from bilateral parietal prior (IC19, Figure 3b) where left ITG activity is not present until 350ms in the no priors approach. Another instance where a constrained solution with fMRI prior seems to elucidate a more detailed pattern of activity is with theta-power approach: while the bilateral frontal network fMRI prior (IC26, Figure 4c) seemed to capture stronger and sustained left orbitofrontal cortex activity, along with less pronounced left ITG contributions compared to no prior approach. The left lateralized fronto-parietal fMRI prior (IC13, Figure 4b) does capture sustained left ITG activity similar to the no prior approach. Additionally, with the occipital prior (IC4, Figure 3c), the early posterior activity is sustained longer than without the prior approach (0-250ms).

### 4.5 A model of dynamic processing across all priors

The event-related active generation fMRI paradigm implicated four broad networks that, when integrated within the MSP framework for MEG source reconstruction, yielded distributed solutions across peristimulus time. Within the first 250ms, solutions constrained to the primary occipital prior (Figure 2c) showed inferior and middle occipital activity within the first 250ms (Figure 3c), followed by left inferior temporal gyrus and orbitofrontal cortex activity (350-500ms). These specific fronto-temporal contributions were apparent within earlier time windows within the MSP reconstruction for the bilateral inferior parietal prior (Figure 2a). With an event-related theta-power approach, the left-lateralized (Figure 2b) and bilateral frontal priors (Figure 2d) displayed sustained inferior temporal gyrus and middle temporal pole (450ms+), along with delayed left OFC activity (750ms+) within the bilateral frontal prior (Figure 4c).

The lack of parietal effects across all MSP reconstructions that is present in fMRI spatial ICA results may suggest parietal contributions may be too scattered temporally to be present in the group analysis, or perhaps that this particular approach may not be sensitive enough to detect these effects within MEG at the group level.

### 4.6 Other considerations

Another important MEG processing consideration that has a potential impact on SNR and interpretability involves treatment of artifacts and characterization of “bad” data. As public sharing of electrophysiological data gains popularity alongside large databases of fMRI data, the MEG/EEG community has continued to build upon methods to improve replicability of experiments. While frequency filtering can address low frequency artifacts, high frequency artifacts like muscle activity, and line noise artifacts, these methods are not able to suppress broadband artifacts and data often require additional inspection. Though many of the widely used software packages (Brainstorm, FieldTrip, MNE, SPM) have ready to use tools that can identify and reject segments of data based on basic metrics, like peak-to-peak signal amplitude differences used here, modern and more advanced methods include identifying bad sensors using Signal Space Separation methods (Jas et al. 2017). There are benefits to automated techniques to identify and reject ‘bad’ data, including reproducibility of data processing across sites and studies, though barriers to successful implementation of these techniques exist, especially when using 4d/BTI MEG data as in the current study.

### 4.7 Conclusions

In summary, the present study built a MEG/fMRI data co-processing pipeline and found four major networks associated with self-generation using fMRI, along with differential progression of activity across multiple sparse priors MEG source reconstruction solutions. Across fMRI priors, evoked responses begin bilaterally across lateral and medial occipital cortex, spreading to inferior frontal, orbitofrontal, and inferior temporal areas. No early significant activity (< 300ms) was captured across analyses examining event-related theta power, and bilateral temporal activity was left lateralized within 750ms. Inclusion of fMRI priors to the MSP framework seemed to produce similar, but more detailed accounts of distributed activity across the cortex.

